# Divergent photosynthesis and leaf respiration responses of four common subtropical tree species to modest warming

**DOI:** 10.1101/2025.05.13.653929

**Authors:** Kashif Hussain, Emily Patience Bakpa, Qiurui Ning, Shihao Huang, Adnan Mustafa, Qianqian Ma, Hui Liu

## Abstract

Subtropical forests are vital to global carbon pools, and their responses to increasing warming may significantly influence carbon sequestration. However, how subtropical tree species adjust the photosynthesis and respiration process in response to climatic warming through phenotypic plasticity is still unclear which is critical for predicting future forest carbon dynamics. In this study we investigate the effects of controlled warming experiment (1.5 ± 0.5) on growth and leaf economic traits especially photosynthesis and respiration in four common subtropical tree species. Our findings reveal that modest warming enhanced photosynthesis, stomatal conductance, and night respiration, leaf anatomical traits as well as growth in *Schima superba, Ormosia pinnata,* and *Pinus massoniana*, of those species under warming. In contrast, *Castanopsis hystrix* demonstrated reduced performance in all measured traits because it may lack adaptive traits critical for warming resilience, indicating a divergent response to the same environmental condition. Our findings highlight the species-specific variation in how subtropical trees adjust traits associated with trees capacity to acclimate to increased temperatures. Specifically, *Ormosia pinnata*, *Pinus massoniana*, and *Schima superba* may exhibit beneficial responses to warming, whereas *Castanopsis hystrix* may experience negative effects in future climatic conditions which alters future carbon storage and tree diversity. Thus, our study offers new insights for further research on common trees under warming to improve predictions of forest dynamics under climate change, aiding conservation efforts to address environmental pressures.

## Introduction

Climate change is exposing organisms to rising temperature and more frequent extreme events (Arnold et al. 2024). Under future scenarios with unmitigated emissions (e.g., SSP5–8.5), the global land surface average temperature is expected to increase by as much as 4.4 by 2100 (Arias et al. 2021; Hazir et al. 2024). High temperature is a universal environmental stressor affecting tree growth and mortality (Wu et al. 2020; Anderegg et al. 2022), influencing forest structure and composition (Brodribb et al. 2020). Plants may possess high natural resilience to warmer temperatures, thereby avoiding thermal stress (Andrew et al. 2023). Alternatively, phenotypic plasticity could enable short-term adjustments to reduce stress exposure or mitigate damage (Brooker et al. 2022). In the tropics, forests dominate the terrestrial carbon cycle, continued increasing temperature may reach critical thermal limits beyond which photosynthetic processes become irreversible (O’Sullivan et al. 2017; Tiwari et al. 2021; Doughty et al. 2023). Subtropical forests also play a role in carbon storage (Li et al. 2019), regulate hydrological cycles and microclimates (Pan et al. 2011; Bordin et al. 2023). Despite beneficial services, global warming poses potential risks to forest structure and functions. Subtropical forests cover about one-quarter of China’s total land area and hold significant ecological importance. A more comprehensive understanding of how subtropical evergreen species in southern China will respond to climate warming and the underlying mechanisms will improve predictions of forest dynamics under future climate conditions.

Photosynthesis and respiration are the two primary biological processes regulating carbon exchange between the atmosphere and the terrestrial biosphere, and the balance between them is important for plant productivity and carbon flux (Aspinwall et al. 2016; Dusenge et al. 2019). The effects of long-term changes in growth temperature on both photosynthesis and respiration are influenced by how these processes acclimate differently to temperature. For example, many species enhance growth by increasing photosynthetic capacity (Drake et al. 2015) more than respiration (Dusenge et al. 2019). The extent to which photosynthesis and respiration acclimate represents a critical factor in elucidating long-term plant responses to environmental change. However, these processes remain inadequately understood for prevalent subtropical species, even with a recent global meta analyses (Wu et al. 2025). Our limited understanding suggests that the degree of acclimation varies between species: fast-growing species tend to acclimate more effectively, while slow-growing species acclimate less. Some species acclimate strongly, while others cannot achieve even partial acclimation (Ow et al. 2008). Elevated temperatures can enhance photosynthesis by improving carboxylation efficiency (Liu et al. 2022), which is achieved through increased concentration and activity of ribulose-1,5-bisphosphate carboxylase/oxygenase (Rubisco) within the optimal temperature range. In contrast, temperatures above the optimum inhibit photosynthesis by disrupting the structure of chloroplasts, damaging the function of photosystem II (Fv/Fm), and reducing the activation state of Rubisco or denaturation of proteins (Didion-Gency et al. 2022), as a result reduction of *A*_sat_ and Fv/Fm was reported due to low soil moisture content (Santos et al. 2018). To the best of our knowledge, there are very few studies which focused on both photosynthesis, respiration and Fv/Fm in response to warming.

In addition, previous studies also focused on structural acclimation and adaptations in response to both short and long term warming. Temperature changes are believed to affect the production capacity, biomass allocation (Wu et al. 2020), and adaptive strategies of plants primarily by altering their functional traits (Zhang et al. 2021). There is contradiction of biomass productivity, some studies showed warming has negative impacts and leads to biomass reduction (Fang et al. 2023), while other studies showed modest warming had positive impacts and increased biomass productivity (Ma et al. 2022; Reich et al. 2022; Liu et al. 2024). Leaf economic traits, such as leaf anatomical traits, that affect tree growth may be key components of adaptive strategies for tree development in the context of climate warming (Slot & Winter 2017; Wu et al. 2018). Chlorophyll (Chl) harvests, transfers, and converts sunlight, making it the key component of photosynthesis. As a result, it is widely recognized as the best indicator of productivity (Li et al. 2018). Terrestrial plants have two types of chlorophyll: Chl A and Chl B, both have strong light absorption capacity and used as indicator of light use efficiency and production efficiency. Previous studies focused on sunlight (Kull et al. 2002), that induces chlorophyll makeup, on the other hand some reported chlorophyll concentration based on taxonomy or functional groups (Halik et al. 2009). However, these conclusions are derived from studies conducted in forested ecosystems or with woody plant species in regions that demonstrate pronounced light gradients between the canopy and understory (Li et al. 2018; Li et al 2018). In contrast, chlorophyll synthesis may also be influenced by temperature. However, how chlorophyll concentration of subtropical forest evergreen species may change under warming has been rarely studied, while grasslands showed divergent responses (Zhang et al. 2021). Furthermore, elevated temperatures can alter stomatal traits, including size, shape, and density (Wu et al., 2018), which in turn affect photosynthesis, respiration and transpiration rates (Hao et al. 2019). However, there are no consistent trends in stomatal plasticity for different tree species under warming conditions (Haworth et al., 2015; Wu et al. 2018; Wu et al. 2020). Increased stomatal density and size induced by warming can enhance the photosynthetic rate (Wu et al. 2018; Pankasem et al. 2024). Previous studies have showed that biomass allocation between above-and belowground, as well as among different organs, is crucial for tree species to acclimate to warming by showing structural and functional adaptations (Wu et al. 2020; Reich et al. 2022). Overall, the plasticity of leaf anatomical traits to warming may facilitate shifts in the photosynthesis, respiration, drought resistance and subsequently tree growth and gross primary productivity.

In this study we conducted a two-year field modest warming of 1.5±0.5 which is predicted to occur in this century to investigate effects of warming on photosynthesis, respiration and leaf economic traits of four co-occuring subtropical evergreen forest species: *Castanopsis hystrix, Ormosia pinnata, Pinus massoniana* and *Schima superba*. These four species are specifically selected due to their common occurrence and distribution range (existence in almost all regions along the altitudinal gradient) from local forests. We quantified the photosynthesis, respiration response, leaf anatomical adaptations, especially how chlorophyll content influences the growth of these species to modest warming. We hypothesized (1) Both photosynthesis and respiration rate would be increased as these species acclimatize to modest warming; (2) modest warming stir growth which is achieved by leaf adaptations especially higher chlorophyll content, and (3) modest warming induces species specific plastic responses that result in divergent adaptive outcomes

## Materials and Methods

### Study site

The study was carried out at South China Botanical Garden in Guangzhou, Guangdong province, China (23°20′ N, 113°30′ E). The climate of the experimental site is subtropical monsoon climate, the average annual temperature is about 22.1L, the coldest month is January, the average temperature is 13.5L, the hottest month is July, the average temperature is 28.6L. The average annual precipitation is 1788.8 mm, the most precipitation month is June, the mean precipitation is 307.4 mm, the least precipitation month is December, the mean amount of precipitation is 29.8 mm. The region has obvious dry and wet seasons, with ∼80% precipitation occur in the wet season from April to September, and others in the dry season from October to March.

### Experimental design and treatment

For the warming experiment, six experimental chmabers were constructed using bricks and concrete. The chambers were 3 m* 3 m in area, 0.8 m in depth, and 0.1 m high above the ground. Three of the six chambers were covered by transparent glass to form hexagonal open top chambers (OTC), and the rest three were used as controls. The total height of the open-top growth box is 3.5 meters, the ground is a regular hexagon with aside length of about 1.2 meters, and the opening part of the top is a regular hexagon with a side length of 0.7 meters. Both control and OTCs chambers receive only ambient precipitations.

The seedlings used in the study were taken from a nursery 2 km away from the study site. All seedlings were grown from the seed of the mother tree, not pruned. At the beginning of the experiment, the seedlings were all one year old. The soil was collected about 30 km away from rural forests, and only a small area of soil was collected to ensure a uniform soil texture. The soil was then naturally air-dried and crushed, and sifted by 2 mm mesh to determine the physiochemical properties of the soil used for seedling cultivation as follow: sand grains 28%, powder grains 46%, mucilage 26%, alkali hydrolyzed nitrogen 16.17 mg kg^-1^, total nitrogen 0.17 g kg^-1^, total phosphorus 0.17 g kg^-1^, total potassium 2.37 g kg^-1^, phosphatase 0.25 mg kg^-1^. One-year old seedlings of *Castanopsis hystrix, Ormosia pinnata, Schima superba* and *Pinus massoniana* were collected, with 10 individuals of each species transplanted into each OTC and control pools on October 2021. All measurements were done after two-year warming treatment.

### Environmental monitoring

Environmental conditions of both control and the OTCs were monitored using a HMP155A temperature probe for air temperature and Campbell 109 thermocouples for soil temperature (5 cm depth). Data were recorded at 15-minute intervals (CR1000, Campbell Scientific, USA) and analyzed for the last two years. We measured all ecophysiological traits during last week of April 2024 to first week of May 2024 (red highlighted part in Fig. 1).

**Fig. 1.**
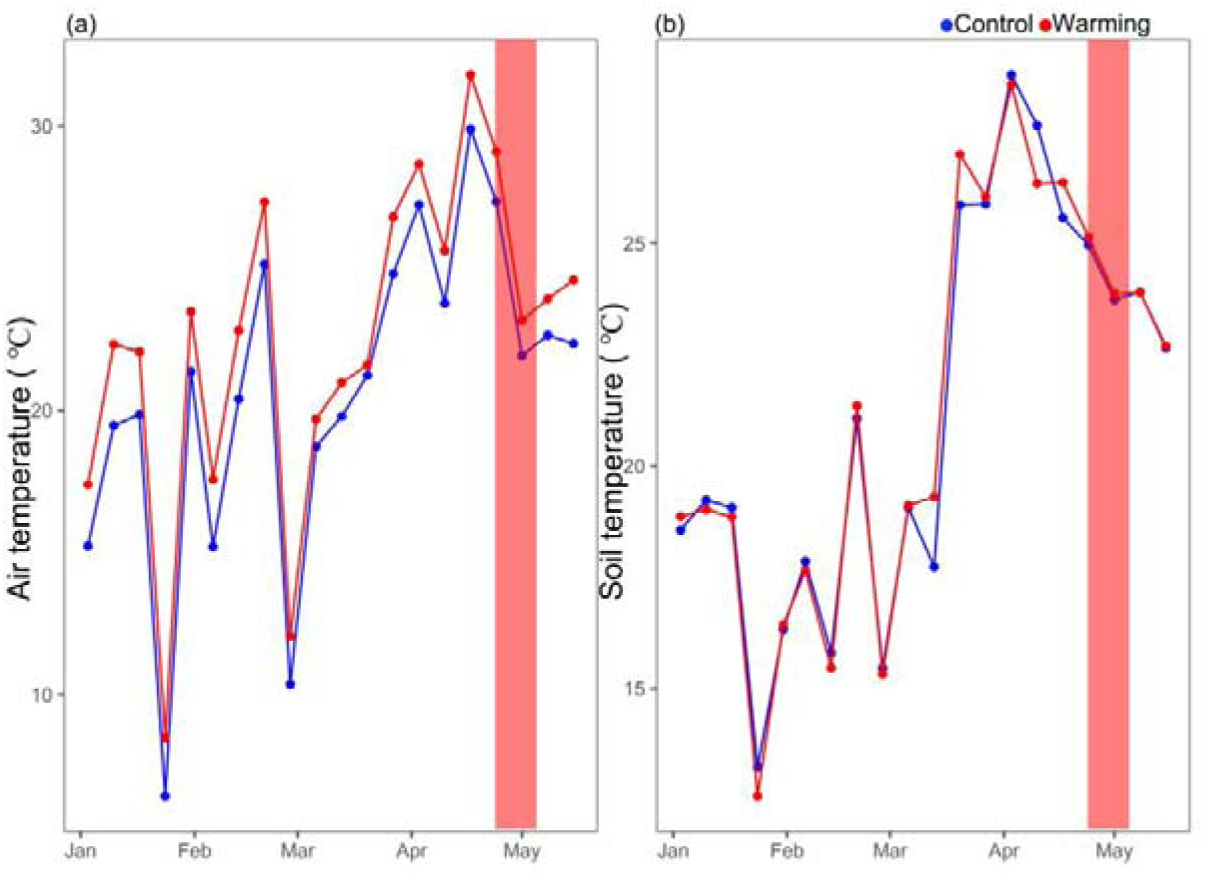
Monthly mean air temperature (**a**) and monthly mean soil temperature (lines) (**b**) of five months at the warming and control plots. Experiment period of this study was highlighted in red.

### Plant Height and Base stem diameter

Plant height and stem diameter were measured for all 10 individuals per species as a vertical distance from the apex of the tree to the level of the soil point where soil-tree contact occurs in both control and OTC at the start of May 2024.

### Leaf gas exchange and chlorophyll fluorescence

After two years of warming treatment, leaf gas exchange was measured from each plot by selecting ten random newly developed fully expanded leaves from five individuals of each species. Photosynthetic rate (A) was measured from 9:00 am to 11:30 am in two consecutive days in the last week of April by using a Li-Cor 6800 infra-red gas analyser equipped with the 2-cm^2^ Multiphase Flash Fluorometer (6800-01A) (Li-Cor BioSciences, Lincoln, NE, USA). Leaves were exposed to a CO_2_ concentration of 400 ppm (using the built-in Li-Cor 6800 CO_2_ mixer), a relative humidity of 50%, and an ambient temperature of 25 L. A was measured at 1500 µmol m^−2^ s^−1^ photosynthetic photon flux density (PPFD), after it had been determined that this light level was saturating, but not damaging, to all species. During measurements with the portable photosynthesis system, five or more needles of *Pinus massoniana* were positioned to fully cover the leaf chamber window. This arrangement was necessary due to the narrow and needle-like structure of *P. massoniana* foliage, ensuring sufficient leaf area coverage for accurate gas exchange readings.

Afterwards, night respiration was measured by using same protocol but keeping PPFD at 0 µmol m^−2^ s^−1^ to fully dark adapted leaves from 02:00 am to 4:00 am at night. To measure chorophyll fluorescence, we again selected fully dark night adapted ten leaves from five individuals of each species by using same Li-Cor 6800 infra-red gas analyser (Li-Cor BioSciences, Lincoln, NE, USA) after stabilized df/dt value near to 0 from 02:00 am to 04:00 am. However, we changed protocol under Fluorometer options by selecting FoFm (dark) or FsFm’ (light), flashtype rectangular, turn light source off, dark mod rate at 50 Hz, red rectangular flash at 8000 µmol m^−2^ s^−1^ for 1000ms duration and output rate at 100 Hz.

To measure photosynthesis temperature (A-T) response curve, leaf photosynthesis was measured at five different temperatures ranging from 15 to 35 °C in 5 °C increments by using Li-Cor 6800. First we measured at 25°C by keeping reference CO_2_ concentration at 400 ppm, relative humidity of 50% and saturating PPFD of 1500 µmol m^−2^ s ^−1^ at 09:00 am. Leaf temperature was controlled using a leaf chamber and measured with an IR sensor. Photosynthetic rates were recorded at temperatures of 10, 15, 20, 25, 30 and 35°C, after a 10-15 minute period at each temperature, afterwards we measured ten leaves from five individual of each species.

### Leaf anatomical traits

Ten fully developed leaves from five individuals of each species of both plots were randomly selected, scanned and the leaf area (LA) measured using LI-3000 Portable Leaf Area Meter (Li-Cor BioSciences, Lincoln, NE, USA). The leaves were then placed in an envelope and oven-dried at 65 for 72 h. The dry mass of the leaves was also measured and the specific leaf area (SLA) equals LA divided by dry mass. To measure stomatal traits, we prepared epidermal peels from leaf surfaces of ten leaves from five individuals per species of each plot. We first applied a layer of transparent nail polish to fix the stomata on the leaves. This nail polish layer was then adhered to a piece of transparent tape cut to 1.0 cm × 1.0 cm. The tape was flattened on a microscope slide and covered with a coverslip for imaging. All images were captured using a microscope (YS100, Nikon, Tokyo, Japan) and analyzed with Image J software. We captured three images of epidermal peels from each of four leaves per species. At 400× magnification, we measured stomatal pore length (SL, μm), and at 100× magnification, we counted the stomata in a 0.25 mm² field of view to determine stomatal density (SD, mm^−2^).

### Leaf chlorophyll contents

Chlorophyll extraction followed Lichtenthaler (1987) with modifications. Ten fresh leaves per species were cleaned, cut into 0.2 g pieces, and homogenized in 5 mL of 80% acetone using a mortar and pestle. The extract was centrifuged at 3000 rpm for 10 min (Centrifuge Model 5425). Absorbance was measured at 663 nm (Chl a), 645 nm (Chl b), and 470 nm (carotenoids) using a UV-Vis spectrophotometer (UV-1000, Shanghai, China). Total chlorophyll content was calculated using the equations of Lichtenthaler (1987):

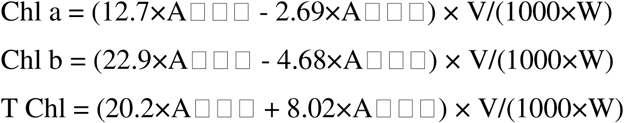

### Leaf water potential

We assessed plant water status by measuring leaf water potential at predawn (Ψpd, MPa) and midday (Ψmd, MPa) using a Scholander-type pressure chamber (PMS 1505D, PMS Instruments, Corvalis, Oregon, USA). Measurements were taken between 05:30 and 07:30 am for predawn and between 13:00 and 14:00 pm for midday (Turner 1988). For each species, we sampled a minimum of ten fully expanded mature leaves from five individuals in both the OTC and control chambers to obtain leaf water potential measurements.

## Statistical analysis

Data were assessed using Kolmogorov–Smirnov test for normality and Levene’s test for homogeneity of variance prior to statistical analysis. Statistical analysis was performed using analysis of variance (ANOVA) followed by the post hoc Tukey HSD test using R software (R core team, 2023). To test if species grown under OTC showed higher response than control plots, traits between control and warming plants were compared using the paired *t*-test (with the *t.test* function in R) and Principal component analysis (PCA) (with the *prcomp* function in R). To test the main effect of warming on each species and their plastic response, we calculated average of each trait of species under control and warming by this formula:

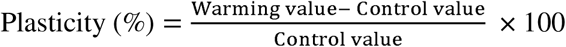

**Table 1.** Summary of variables measured in this study.

| Traits | Description | Unit |
| --- | --- | --- |
| PH | Plant height | m |
| BD | Base stem diameter | cm |
| Chl A | Chlorophyll A concentration | mg g<sup>-1</sup> |
| Chl B | Chlorophyll B concentration | mg g<sup>-1</sup> |
| T Chl | Total chlorophyll concentration | mg g<sup>-1</sup> |
| Carotenoids | Total carotenoids | mg g<sup>-1</sup> |
| A | Photosynthetic rate | μmol m<sup>-2</sup> s<sup>-1</sup> |
| g<sub>s</sub> | Stomatal conductance | mmol m<sup>-2</sup> s<sup>-1</sup> |
| R<sub>n</sub> | Night respiration | μmol m<sup>-1</sup> s<sup>-1</sup> |
| Fv/Fm | PSII efficiency | unitless |
| SL | Stomatal length | μm |
| SD | Stomatal density | mm<sup>-2</sup> |
| LA | Leaf area | cm<sup>2</sup> |
| SLA | Specific leaf area | cm<sup>2</sup> g<sup>-1</sup> |
| Ψ<sub>pd</sub> | Predawn water potential | MPa |
| Ψ<sub>md</sub> | Midday water potential | MPa |

## Results

### Environmental variables

During the last five months from Jan 2024 to first week of May 2024 the average monthly air temperature of OTC was significantly higher by 1.5 ± 0.5L compared to the temperature of controlled plots (Fig. 1a, *p* = 0.31). The average monthly soil temperature of OTC was higher by 0.11L compared to the soil temperature of controlled plots (Fig. 1b, *p* = 0.50).

### Leaf gas exchange and chlorophyll fluorescence

A and g_s_ showed significant positive responses in *O. pinnata*, *P. massoniana* and *S. superba* while both A and g_s_ response of *C. hystrix* under warming decreased significantly (mean plasticity values of A and g_s_ are 19% and 18%, respectively, Fig. 2 a, b; Table S1). Night respiration in all species showed significant positive response to warming as compared to control plots, however, Fv/Fm showed negative response under warming in all species (mean plasticity is 30% and -2.4% for Rn and Fv/Fm, respectively, Fig. 2 c, d; Table S1).

**Fig. 2.**
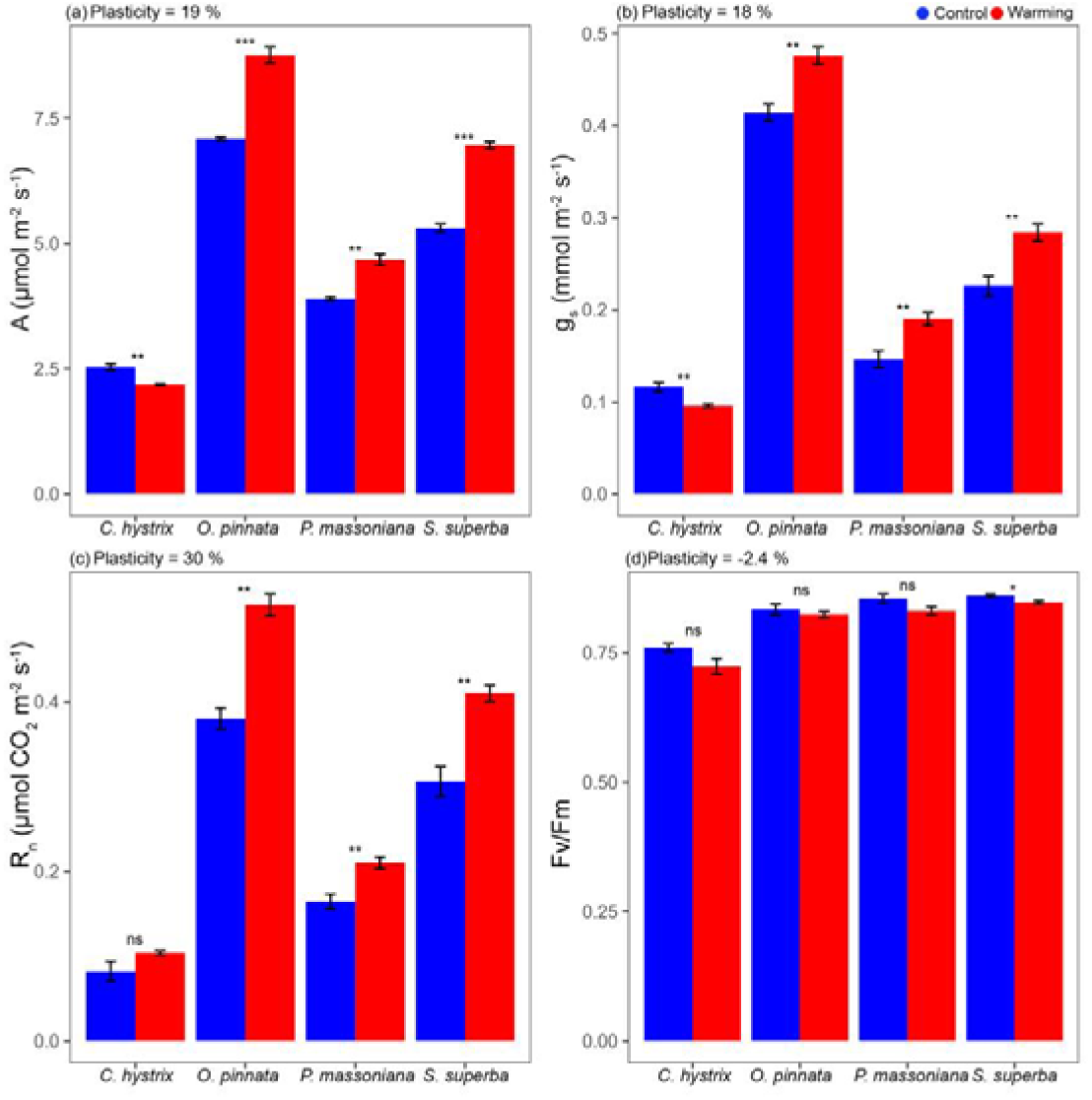
Photosynthetic rate (**a**), stomatal conductance (**b**), night respiration (**c**) and photosystem II efficiency (Fv/Fm, **d**) for *Castanopsis hystrix*, *Ormosia Pinnata, Pinus massoniana and Schima superba* grown in the control and warming site . Error bars are standard error (n = 5). Asterisks (*), (**) and (***) indicate that there are significant differences at *p* < 0.05, *p* < 0.01 and *p* < 0.001 between the control and warming plots, respectively. Mean values of plasticity of the four species are reported for each trait.

The temperature response curves followed a parabollic shape for all species under both conditions. *C. hystrix* showed increasing trend of A response but lower than control plot individuals (Fig. 3, a). However, *O. pinnata*, *P. massoniana* and *S. superba* showed higher response to temperature which showed their T _opt_ lies between 25-30 (Fig. 3 b,c,d).

**Fig. 3.**
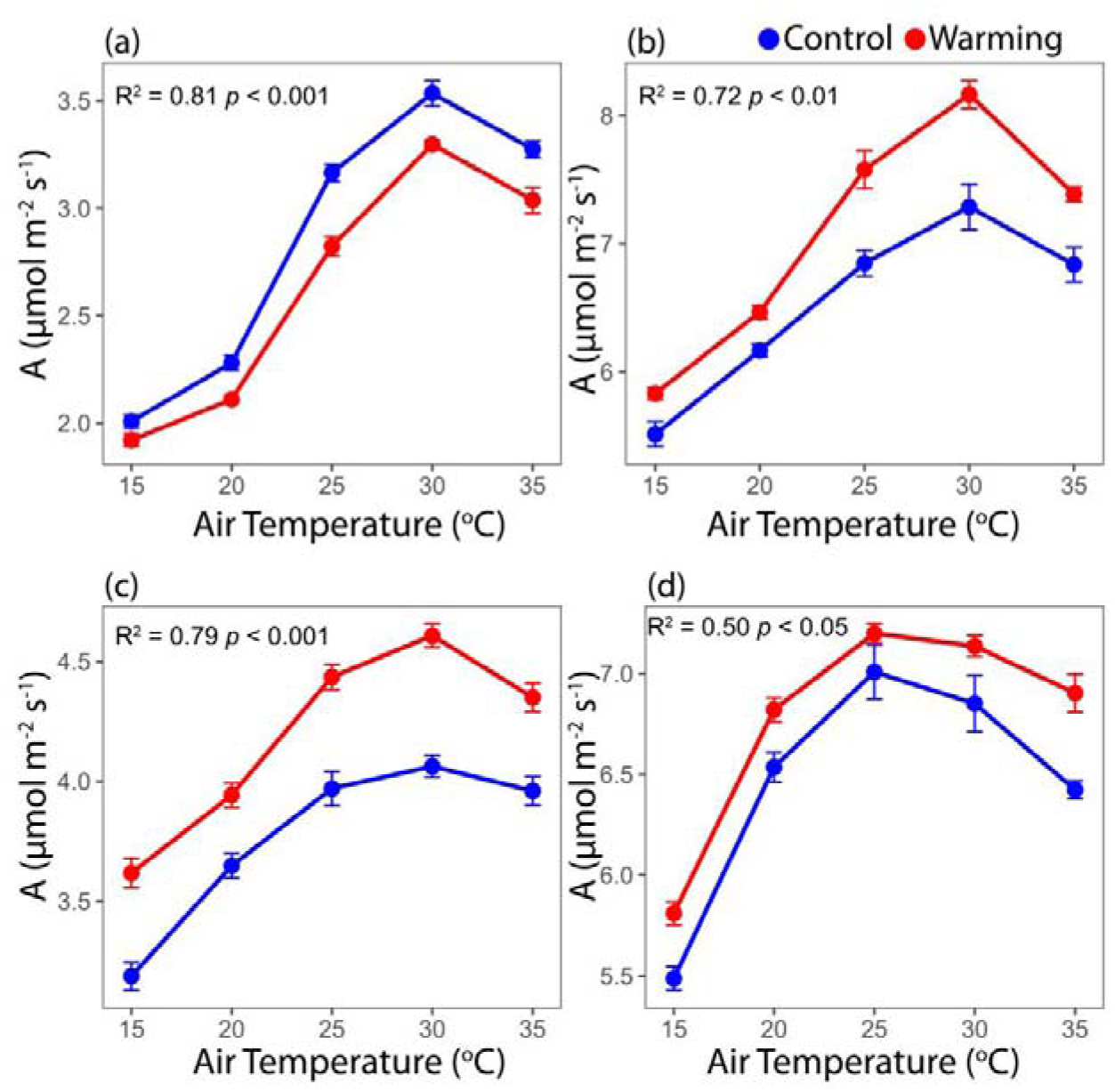
Temperature response curves of photosynthesis (A) of *Castanopsis hystrix*, *Ormosia Pinnata, Pinus massoniana and Schima superba* grown in the warming site and control site. The points represent the absolute net photosynthetic rate at each temperature.

### Plant height and DBH

Under two years of warming, plant height increased significantly of *O. pinnata*, *P. massoniana* and *S. superba*, while we saw a slight decrease in plant height of *C. hystrix* as compared to controlled plots (mean plasticity value of plant height 13.8%, Fig. 4a, Table S1). DBH response of *O. pinnata* showed significant positive response to warming, while other three species showed no significant differences in DBH (mean plasticity value of DBH 14.5%, Fig. 4b, Table S1).

**Fig. 4.**
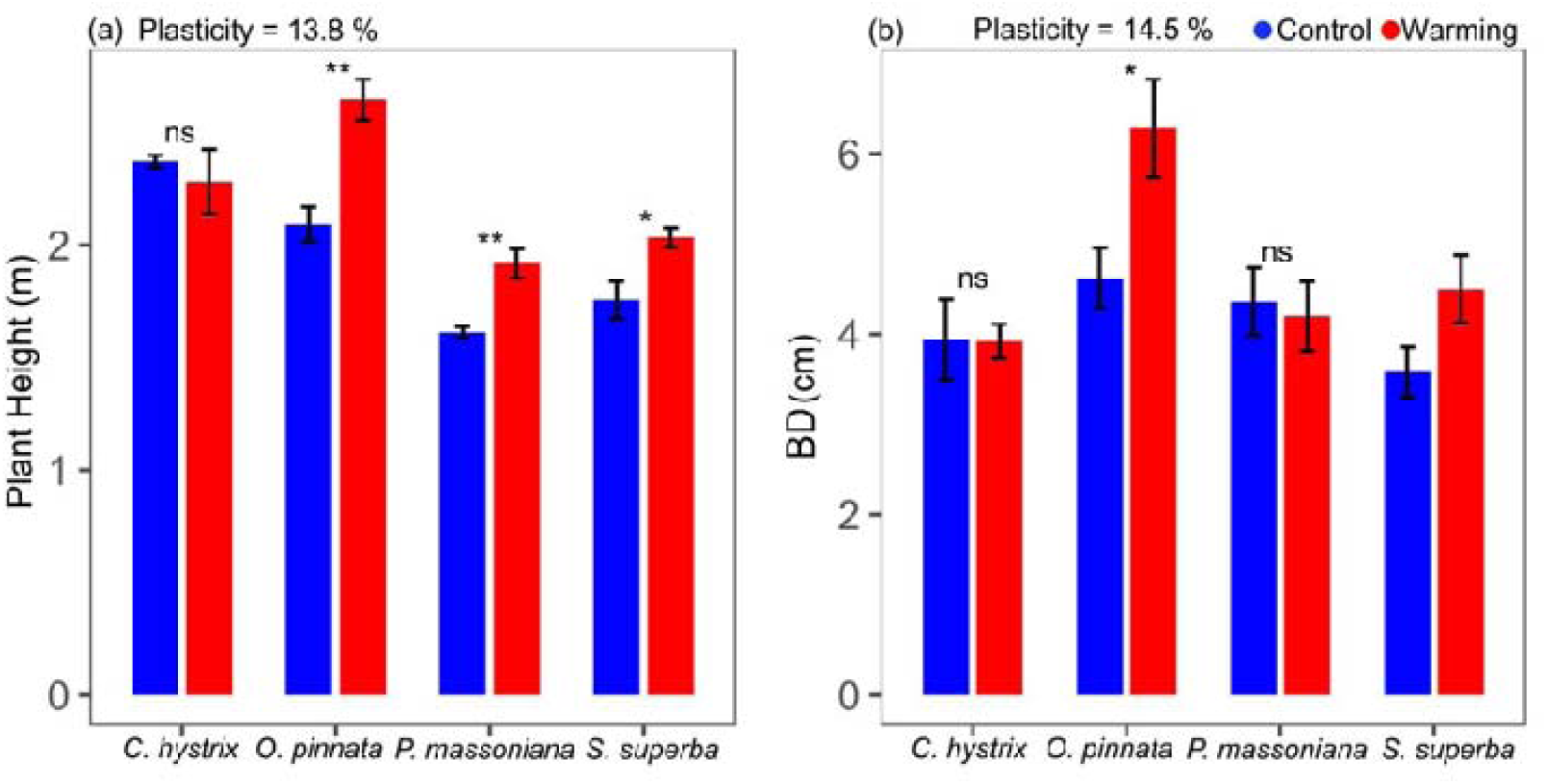
The Plant height (**a**), the stem diameter (**b**, BD) for *Castanopsis hystrix*, *Ormosia Pinnata, Pinus massoniana* and *Schima superba* grown in the warming site and control site. Error bars are standard error (n= 5). Asterisks (*) and (**) indicate that there are significant differences at *p*< 0.05 and *p*< 0.01 between the warming and the control plots respectively. Mean values of plasticity of the four species are reported for each trait.

### Leaf structural traits

Overall, leaf area and specific leaf area of all species under warming increased significantly (mean plasticity is 24.7% and 10.6% for LA and SLA, respectively, Fig. S2 a, b, Table S1). In addition, both stomata length and density increased significantly in response to warming of *O. pinnata*, *P. massoniana* and *S. superba*, while both length and density decreased significantly in *C. hystrix* under warming (mean plasticity is 8.4% and 7.7% for SL and SD, Fig S2 c, d, Table S1). Furthermore, all photosynthetic pigments showed positive responses to warming (Tabel S1). Specifically Chl A, increased significantly in *P. massoniana*, while Chl B increased significantly in all species (mean plasticity is 11.6% and 41.8% for Chl A and Chl B, Fig S3 a, b, Table S1). T Chl and carotenoids also showed higher trend in response to warming (mean plasticity is 20.7% and 28% for T Chl and Carotene, Fig S3 c, d, Table S1)

### Leaf water potential

Both predawn and midday water potentials under warming became more negative. *C. hystrix* was more negative under warming in response to control than other species (Fig S4, Table S1). Afterwards, *P. massoniana* then *O. pinnata* and lastly *S. superba* had significant negative values for both predawn and midday water potentials (mean plasticity is 15.8% and 9% for Ψpd and Ψmd respectively, Fig S4, Table S1).

### Correlation between physiological and structural traits

Photosynthetic rate and night respiration showed positive correlations with stomatal conductance among all species (Fig. 5 a,b), with the non-significant correlation existed between night respiration and stomatal conductance for *P. massoniana* and *C. hystrix* (Fig. 5, b *p* = 0.05, *p* = 0.16). On the other hand, photosynthetic rate showed significant correlations with leaf area for three species except *P. massoniana* (Fig. 5 c), and night respiration only showed significant correlation with leaf area for *O. pinnata* (Fig. 5 d). However, night respiration of *C. hystrix* showed a significant negative correlation with gs and also a significant negative correlation of photosynthetic rate with leaf area (Fig. 5 b,c). Both photosynthesis and night respiration showed positive correlations with stomata length (Fig S5 a,b) and stomata density (Fig S5 c,d).

**Fig. 5.**
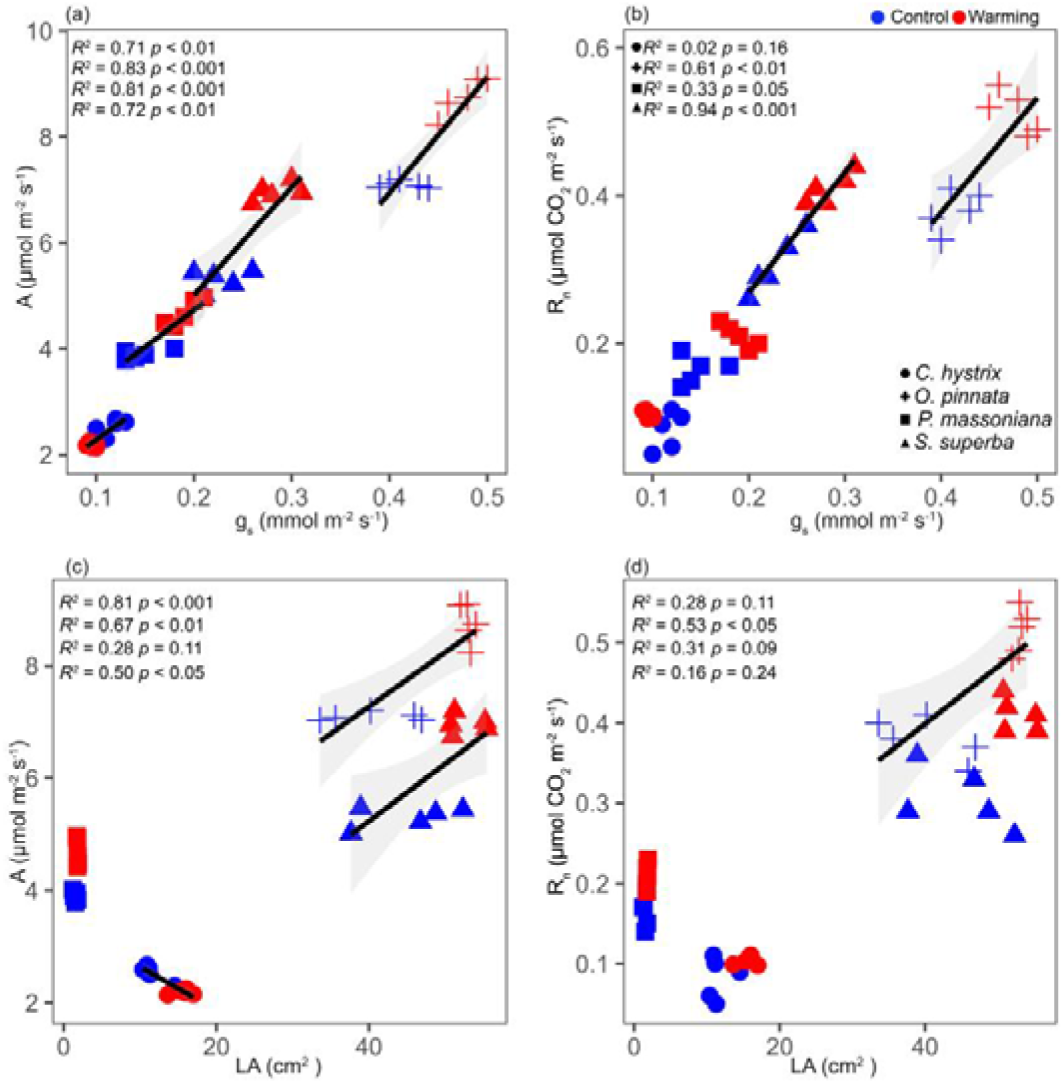
The relationships between A and gs (a), night respiration (R_n_) and g_s_ (b), A and R_n_ with LA (c), (d). Species under control are shown in blue; Warming condition species in red. Only statistically significant relationships (*p*<L0.05) are shown as solid lines with shaded areas representing the 95% confidence interval.

### Principal component analyses

According to the PCA results, control and warming plot plants were separated in the space defined by the first two principal components (PCs). PC1 and PC2 explained 52.5% and 17.4% of the total variation, respectively (Fig. 6). Growth related traits clustered together on the negative side of PC1, whereas Ψpd and Ψmd directed towards positive side of PC1. Most of the leaf physiological and anatomical traits located on the negative side of PC2, leading by A, g_s_, R_n_, SD and SL. Overall PC1 represents that warming influences traits related to growth, potentially indicating an enhancement in these traits under warmer conditions, while PC2 indicates physiological stress response differences between the treatments. Species under warming showed higher growth (height and SLA) and lower Fv/Fm (Fig. 6).

**Fig. 6.**
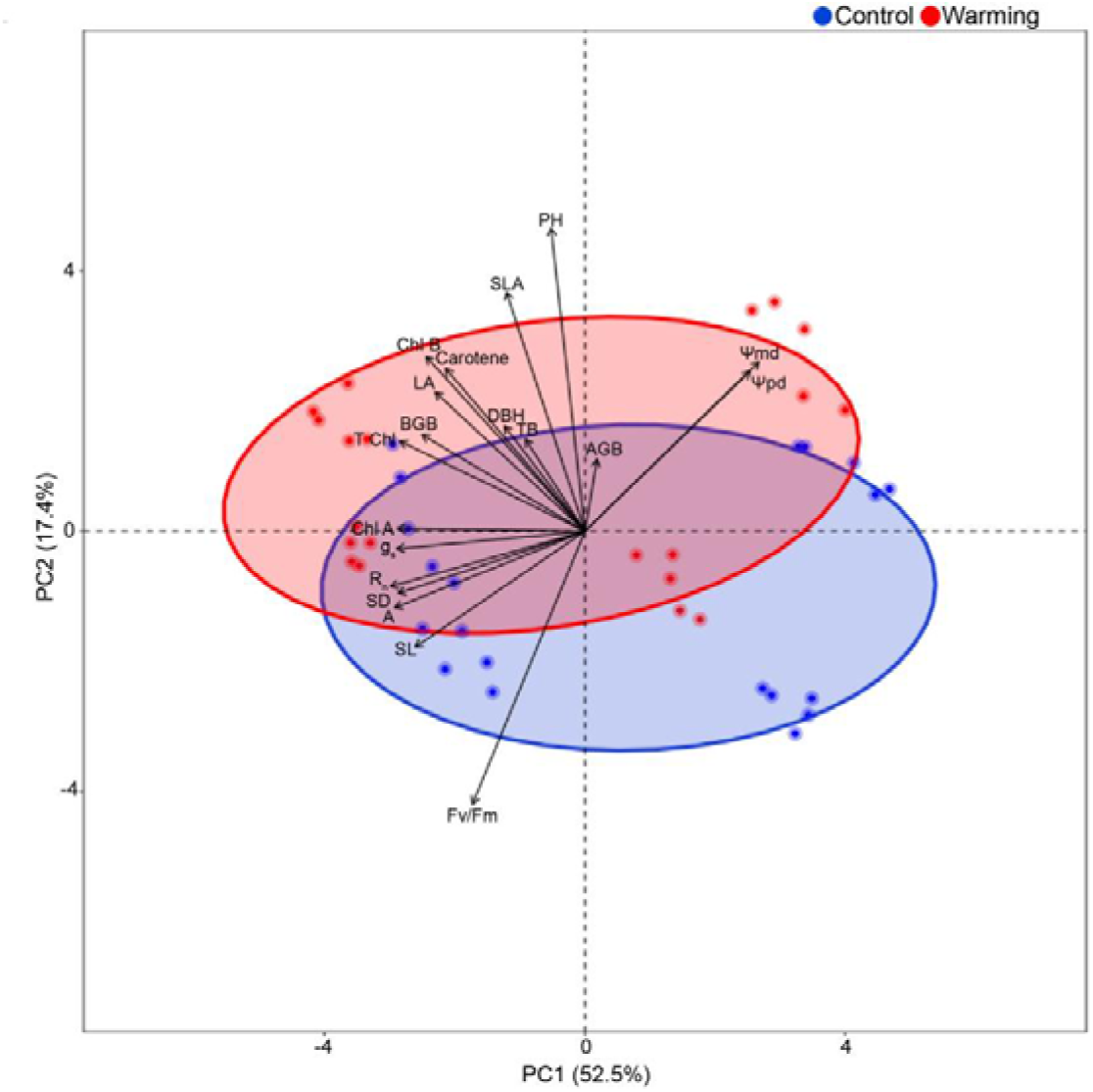
Principal component analysis of control and warming plot species based on 19 leaf functional traits in this study. Control species are shown in blue, warming species are shown in red.

## Discussion

In this two-year OTC warming study, we found that (1) warming overall increased photosynthesis, respiration response of subtropical forest species and decreased Fv/Fm of all species which support our hypothesis and supported by previous studies (Slot & Kitajima 2015; Slot & Winter, 2017; Travinen et al. 2022) (Fig. 2, Fig. 7), and (2) increasing trend of height, DBH and leaf structural traits were observed under warming and also water potential tended to be more negative as compared to control plots also support our hypothesis and (Wu et al. 2020) reported increased DBH (Fig. 3, Fig. S1, S2). (3) We found three species *O. pinnata, P. massoniana* and *S. superba* acclimated well to warming by showing significant plastic responses to all studied traits in this study except *C. hystrix*. Our species specific responses are supported by a recent study (Crous et al. 2025; Wu et al. 2020) they also reported reduced photosynthetic rate, increased dark respiration to warming in tropical species and subtropical specific species showed divergent plastic responses to warming. Warming increased photosynthetic rate and night respiration for *O. pinnata, P. massoniana* and *S. superba* which were correlated with higher g_s_, leaf area and stomatal traits, thus these species grow rapid and accumulate more biomass. All species increased chlorophyll and carotenoid pigments significantly under warming. Interestingly, *C. hystrix* under warming showed negative growth, plant height, basal diameter, photosynthetic response, stomatal traits but increased leaf area and night respiration. The decline in growth for *C. hystrix* may be due to lower photosynthetic rate, correlating with stomatal conductance and stomatal density. Overall, our study offers new insights into how subtropical forest species acclimatize to modest warming by increasing photosynthesis, respiration and declining Fv/Fm, highlighting a metabolic adjustment mechanism (Wahid et al. 2007). Our study reveals species-specific responses to modest warming in subtropical forests, with critical implications for forest management under climate change. Enhanced photorespiration and growth in *Ormosia pinnata*, *Pinus massoniana*, and *Schima superba* suggest these species are resilient to moderate temperature increases and could be prioritized in reforestation and afforestation programs. In contrast, *Castanopsis hystrix* exhibited reduced performance across all measured traits, indicating its vulnerability to warming. This divergence underscores the need for adaptive management strategies to maintain ecosystem stability and carbon sequestration in subtropical forests.

**Fig. 7.**
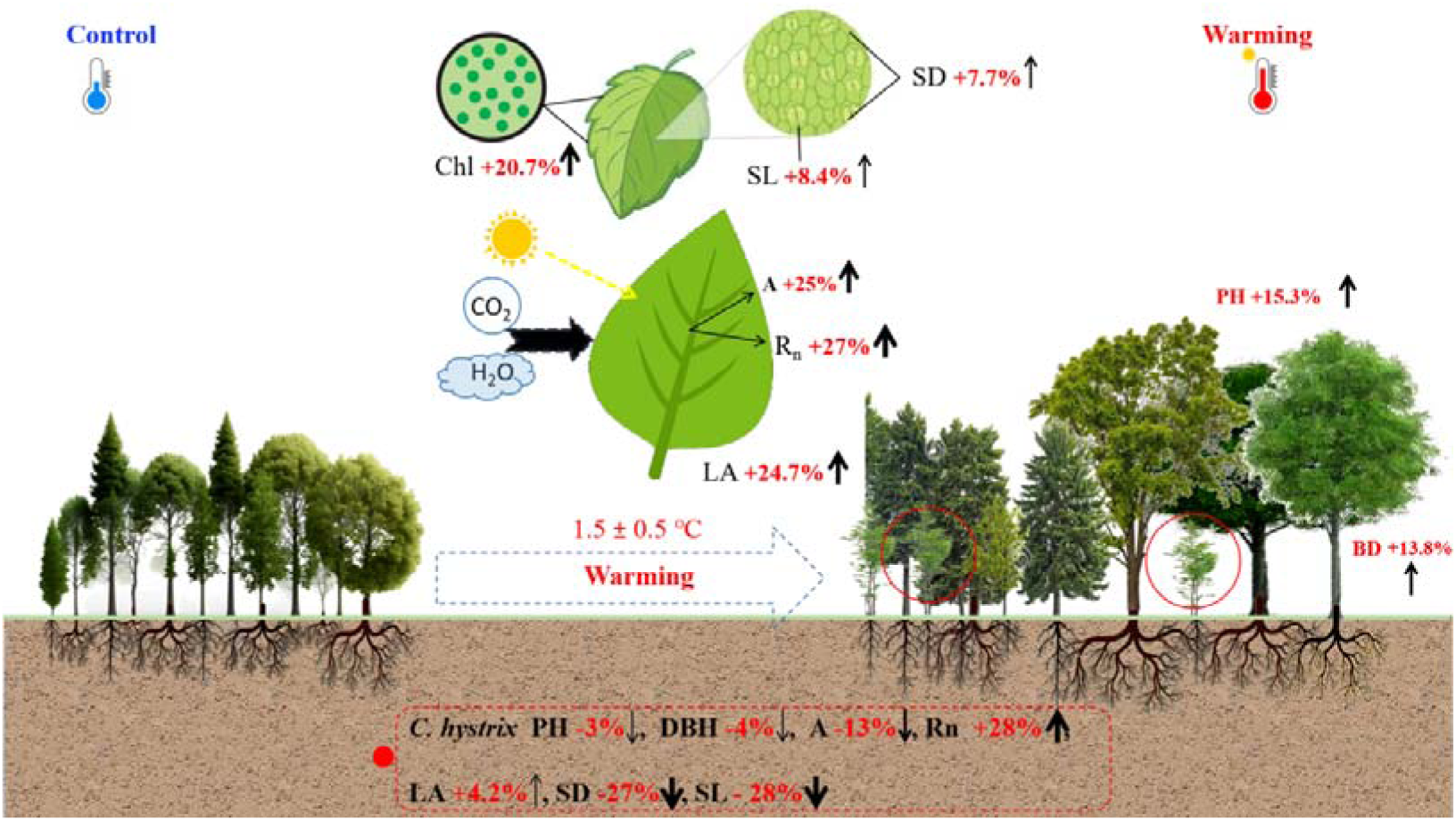
A schematic diagram showing the influence of modest warming on four common subtropical tree species. Data are averaged from those under control and warming treatment. The up or down arrows and width indicate a significant increasing or decreasing trend respectively. *C. hystrix* divergent response showed in red circle and dotted red rectangle.

### Acclimation of photosynthesis and respiration to warming

Warming exhibited contrasting effects depending on species. Warming benefited photosynthetic rate, stomatal conductance of *O. pinnata*, *P. massoniana* and *S. superba*. However, photosynthetic rate and stomata conductance of *C. hystrix* was negatively impacted by warming of 1.5 ± 0.5L. Biochemical limitations may have partially contributed to this effect, as evidenced by the observed decrease in maximum Fv/Fm, low A-T response, and the positive correlations among A-gs, A-LA, A-SL, and A-SD (Fig. 5, Fig. S5). Our results for *C. hystrix* under warming were supported by previous study reported lower assimilation rate in beech plant and tropical mature trees due to severe stomatal limitation and reduced stomatal conductance (Didion-Gency et al. 2022; Crous et al. 2025). On the contrary, *O. pinnata*, *P. massoniana* and *S. superba* showed higher stomatal density and length, as well as increased photosynthesis and stomatal conductance in three species (Wu et al. 2020). In addition, these three species showed better performance because they acclimatize to modest warming which showed contradiction to a recent study by (Crous et al. 2025), which reported downregulation of photosynthesis and respiration capacity of tropical trees. We found all species decreased Fv/Fm in response to warming which were due to partial inhibition of PSII, an increase in energy dissipation as heat and photoinhibition of the photosynthetic apparatus. The same decreased results were reported by (Guchou et al. 2007) for *C. hystrix* and *S. superba* under modest warming.

Overall, we found increased night respiration in all species under warming (Fig. 2). Night respiration showed significant positive trends with stomatal conductance, leaf area, stomata length and density (Fig. 5, Fig. S5) except for *C. hystrix* and *Pinus massoniana*. This was supported by Li et al. (2017) who reported leaf respiration acclimated more strongly than photosynthesis in response to warming, which led to an increase in carbon assimilation while moderating carbon losses. In addition, higher night respiration of *C. hystrix* induced by warming may be due to increased mitochondrial activity (Scafaro et al. 2021) and increased maintenance cost raises respiration independent of daytime photosynthesis (Atkin et al. 2003) supporting the idea that warming directly elevates respiration. Our findings differed from several prior studies on tropical trees, which indicated a reduction in respiration as a respiratory thermal acclimation response to increasing temperatures (Slot & Winter, 2017; Crous et al. 2025). Wang et al. (2020) proposed an optimality theory of photosynthetic capacity, which demonstrated a close link between respiratory thermal acclimation and the thermal acclimation of photosynthetic capacity. This relationship ensures optimal photosynthesis and efficient resource use. Therefore, under modest warming, both Rubisco and mitochondrial enzymes trigger the Calvin cycle, electron transport chain which increased photosynthesis and respiration. In addition, likelihood of enhanced coordination between carbon fixation for carbohydrates formation and carbon use (night respiration) utilize for growth and maintenance (Dusenge et al. 2019). These results highlight species-specific acclimation strategies, with three species using structural and physiological plasticity to thrive under warming, while *C. hystrix* struggled to cope with thermal stress.

### Warming effects on plant growth

Modest warming led to increased plant growth in *O. pinnata*, *P. massoniana* and *S. superba* due to higher photosynthetic rate; however, a decline in growth was observed in *C. hystrix* (Fig. 3). It has been proposed that increased temperatures may enhance the growth of herbaceous monocots, woody gymnosperms, and species of eucalyptus (Li et al. 2017). Similar to our results, previous studies also found that warming increased height and biomass of *S. superba*, *P. massoniana* and non-significant effect on growth of *C. hystrix* (Li et al. 2017; Wu et al. 2020). Thus, the elevated photosynthetic rates observed in *O. pinnata*, *P. massoniana*, and *S. superba* in our study may have contributed to the enhancement of their height. However, opposite to increasing trend, *C. hystrix* showed decrease in plant height, basal diameter and total biomass (Fig. 3, Fig. S1). The reduction in photosynthetic rate, stomatal conductance (Fig. 2) may have limit the growth of *C. hystrix* which shared same non-significant decreased response of *M. breviflora* under warming (Wu et al. 2020). Our study’s growth results contradict findings that tropical and subtropical tree species may be nearing a high-temperature threshold (Bowman et al. 2014). This discrepancy may be attributed to variations in plant adaptability and local soil water conditions in a warmer environment (Doughty et al. 2023).

### Warming effects on leaf structural traits and photosynthetic pigments

Numerous structural adjustments may have facilitated plant acclimation to elevated temperatures. In this study warming significantly increased leaf area and specific leaf area of all species except a non-significant increasing trend in *P. massoniana*, which indicates that there is a greater amount of leaf area displayed per unit mass in these four species (Li et al. 2017; Liu et al. 2023) (Fig. S2). Increasing trend of SLA linked to more efficient light capture, which may result in greater assimilation gains (Zhu et al. 2020). We found a significant positive correlation between photosynthetic rate and leaf area of all species except *P. massoniana* (Fig. 5) and this is due to increasing leaf area provides extra space to capture more light for photosynthesis. In addition to SLA, an increased photosynthetic rate under warming conditions may be linked to warming-induced changes in stomatal density and length (Fig. S2). These two traits serve as primary regulators influencing the response of stomatal conductance to climatic stresses (Franks et al 2015), consequently, variations in stomatal conductance impact the rate of photosynthesis. We found an increasing trend in both stomata length and stomata density in all species except *C. hystrix* showing decreased both length and density. Our results for *O. pinnata*, *P. massoniana* and *S. superba* showed contradiction with prior studies while *C. hystrix* showed resemblance (Wu et al. 2018; Wu et al. 2020). Lower stomata density can help prevent excessive water use in plants, making it a potential way for enhancing drought tolerance (Franks et al. 2015), which suggests monitoring stomatal traits as early indicators of climate stress.

Overall, we found an increasing trend of photosynthetic pigments in all species in response to warming. The majority of energy utilized for photosynthesis in plants is obtained from light energy captured by photosynthetic pigments. Consequently, the chlorophyll content is intricately associated with the photosynthetic capacity of plants (Zhu et al. 2020). Our results are consistent with prior studies which showed warming enhances the photosynthetic capacity of plants by accelerating the synthesis of photosynthetic pigments, thereby, promoting growth (Sun et al. 2015; Zhu et al. 2020). Meanwhile, warming of 1-2 provides optimal temperature environment for the synthesis of photosynthetic pigments which enhances photosynthetic capacity. Finally, all species under warming showed more negative predawn and midday water potentials, which may be due to adjustments in stomata as well as more soil water consumption and evaporation. Specifically, *C. hystrix* and showed more negative water potential which reduced photosynthetic rate and stomatal conductance under warming, as compared to *O. pinnata* and *S. superba*. These results showed resemblance with a recent study which showed *Q. coccifera, Q. ilex* reduced A net and gs with decreasing water potential (Gauthey et al. 2024).

## Conclusions

Our two year OTC experiment of co-occuring subtropical forest species sheds light on the effect of modest warming on the acclimation of photosynthesis, respiration and growth. Compared to the control, three species under warming increased photosynthesis, respiration and growth except *C. hystrix*. *C. hystrix* under warming decreased photosynthesis but increased night respiration and also decreased total biomass and leaf stomatal traits while other three species showed increased height, basal diameter photosynthesis and respiration. This suggests that subtropical species acclimatize their photosynthesis and respiration differently, which may affect species diversity because only the best-adapted species thrive well in response to future warming. It also suggests actionable strategies for subtropical forest management under climate change, prioritizing resilient species in reforestation to sustain productivity and carbon storage while implementing conservation measures for vulnerable species by integrating physiological monitoring to detect early stress and adopting adaptive practices like mixed-species plantings to maintain biodiversity.

## Acknowledgements

This work was supported by Guangdong Basic and Applied Basic Research Foundation (2024B1515020067), National Natural Science Foundation of China (32371575, 32371641), Guangdong Science and Technology Plan Project (2023B1212060046), the Youth Innovation Promotion Association of the Chinese Academy of Sciences (Y2023093, 2023363, Y202077), and Guangzhou Municipal Science and Technology (2024A04J6433).

## Author contributions

Hui Liu and Qianqian Ma designed the research. Kashif Hussain performed experiments and conducted fieldwork. Kashif Hussain performed analyses with guidance from Hui Liu. Kashif Hussain wrote the first draft of the manuscript. Kashif Hussain, Qiuri Ning, Emily Patience Bakpa, Adnan Mustafa and Hui Liu revised the manuscript.

## Data availability

The data that support the results of this study are available from the corresponding author upon reasonable request.

## Supplementary data

**Table. S1.** Summary of all traits measured in this study for species growth in the warm and control site. Data are means± SE. t value with significant stars.

| Classification | Traits | C. Hystrix Mean± SE |  |  | O. Pinnata Mean± SE |  |  | P. Massoniana Mean± SE |  |  | S. Superba Mean± SE |  |  |
| --- | --- | --- | --- | --- | --- | --- | --- | --- | --- | --- | --- | --- | --- |
|  |  | Control | Warming | *t* value | Control | Warming | *t* value | Control | Warming | *t* value | Control | Warming | *t* value |
| Growth | PH | 2.3 ± 0.02 | 2.2 ± 0.1 | **0.61** | 2.09 ± 0.07 | 2.65 ± 0.09 | **-4.66\*\*** | 1.6 ± 0.02 | 1.9 ± 0.0 | **-4.35\*\*** | 1.76 ± 0.08 | 2.03 ± 0.04 | **-2.94\*** |
|  | DBH | 3.94 ± 0.4 | 3.92 ± 0.1 | 0.04 | 4.6 ± 0.3 | 6.2 ± 0.5 | **-2.59\*** | 4.3 ± 0.37 | 4.3 ± 0.33 | 0.29 | 3.58 ± 0.2 | 4.5 ± 0.3 | -1.96 |
|  | TB | 307 ± 26 | 298 ± 17.5 | **0.27** | 338 ± 20.6 | 457 ± 23.3 | **-3.57\*** | 328 ± 27.7 | 374 ± 27.4 | -1.2 | 264 ± 17.6 | 330 ± 10.2 | **-3.2\*** |
| Physiology | A | 2.5 ± 0.06 | 2.1 ± 0.02 | **5.13\*\*** | 7.06 ± 0.03 | 8.76 ± 0.1 | **-10.2\*\*\*** | 3.9 ± 0.03 | 4.6 ± 0.1 | **-6.77\*\*** | 5.31 ± 0.08 | 6.96 ± 0.07 | **-15.8\*\*\*** |
|  | gs | 0.11 ± 0.005 | 0.09 ± 0.001 | **3.73\*** | 0.41 ± 0.009 | 0.47 ± 0.009 | **-4.72\*\*** | 0.14 ± 0.009 | 0.19 ± 0.007 | **-3.77\*\*** | 0.22 ± 0.01 | 0.28 ± 0.009 | **-4.08\*\*** |
|  | Rn | 0.08 ± 0.01 | 0.1 ± 0.002 | -1.84 | 0.38 ± 0.01 | 0.51 ± 0.01 | **-6.06\*\*\*** | 0.16 ± 0.008 | 0.21 ± 0.007 | **-4.1\*\*** | 0.30 ± 0.01 | 0.41 ± 0.009 | **-5.22\*\*** |
|  | Fv/Fm | 0.75 ± 0.08 | 0.72 ± 0.01 | 2.15 | 0.83 ± 0.01 | 0.82 ± 0.005 | 2.1 | 0.85 ± 0.008 | 0.83 ± 0.007 | **2.1** | 0.86 ± 0.002 | 0.84 ± 0.003 | **2.96\*** |
| Anatomy | CHL A | 0.66 ± 0.05 | 0.79 ± 0.03 | -2.08 | 1.07 ± 0.01 | 1.08 ± 0.005 | -1.11 | 0.76 ± 0.01 | 0.92 ± 0.005 | **-6.81\*\*** | 0.97 ± 0.03 | 1.05 ± 0.02 | -2.56 |
|  | CHL B | 0.35 ± 0.02 | 0.57 ± 0.03 | **-5.07\*\*** | 0.59 ± 0.03 | 0.71 ± 0.003 | **-3.52\*** | 0.34 ± 0.003 | 0.4 ± 0.002 | **-15.6\*\*\*** | 0.46 ± 0.01 | 0.76 ± 0.02 | **-10.6\*\*\*** |
|  | T CHL | 1.02 ± 0.06 | 1.37 ± 0.07 | **-3.5\*\*** | 1.6 ± 0.04 | 1.8 ± 0.007 | **-3.15\*** | 1.1 ± 0.008 | 1.3 ± 0.005 | **-18.18\*\*\*** | 1.44 ± 0.04 | 1.82 ± 0.02 | **-7.7\*\*\*** |
|  | Carotene | 0.6 ± 0.04 | 1 ± 0.02 | **-7.25\*\*\*** | 1.02 ± 0.01 | 1.07 ± 0.004 | **-2.65\*** | 0.52 ± 0.01 | 0.95 ± 0.01 | **-9.09\*\*\*** | 0.94 ± 0.04 | 0.97 ± 0.02 | **-0.67** |
|  | **SD** | 16.2 ± 0.3 | 11.8 ± 0.2 | **10.37\*\*\*** | 23.2 ± 0.5 | 28.2 ± 0.3 | **-7.21\*\*\*** | 15 ± 0 | 15.6 ± 0.24 | -2.49 | 28.4 ± 0.8 | 33.8 ± 0.5 | **-5.14\*\*** |
|  | **SL** | 17.6 ± 0.3 | 12.7 ± 0.2 | **9.45\*\*\*** | 21.5 ± 0.4 | 25.2 ± 0.2 | **-7.39\*\*\*** | 16.3 ± 0.5 | 21.4 ± 0.21 | **-8.09\*\*\*** | 30.2 ± 0.2 | 33.4 ± 0.4 | **-5.98\*\*** |
|  | **LA** | 11.6 ± 0.7 | 15.6 ± 0.5 | **-4.26\*\*** | 40.4 ± 2.6 | 53 ± 0.3 | **-4.76\*\*** | 1.51 ± 0.1 | 1.8 ± 0.02 | -2.74 | 44.8 ± 2.8 | 52.7 ± 1.07 | **-3.8\*** |
|  | **SLA** | 14.7 ± 0.7 | 15.3 ± 0.2 | -0.81 | 12.7 ± 0.2 | 14.8 ± 0.3 | **-4.4\*\*** | 1.9 ± 0.09 | 2.8 ± 0.3 | -2.42 | 11.9 ± 0.18 | 12.7 ± 0.16 | **-3.09\*** |
| **Water** | **Ψ<sub>pd</sub>** | -0.94 ± 0.009 | -1.04 ± 0.01 | **-7.01\*\*\*** | -0.49 ± 0.01 | -0.62 ± 0.01 | **-8.09\*\*\*** | -0.74 ± 0.009 | -0.84 ± 0.01 | **-6.38\*\*\*** | -0.35 ± 0.01 | -0.44 ± 0.007 | **-6.6\*\*\*** |
|  | **Ψ<sub>md</sub>** | -1.71 ± 0.01 | -1.89 ± 0.01 | **-10.7\*\*\*** | -0.83 ± 0.01 | -0.96 ± 0.01 | **-7.15\*\*\*** | -1.14 ± 0.01 | -1.38 ± 0.01 | **-10.7\*\*\*** | -0.74 ± 0.008 | -0.85 ± 0.01 | **-7.90\*\*\*** |

**Fig. S1.**
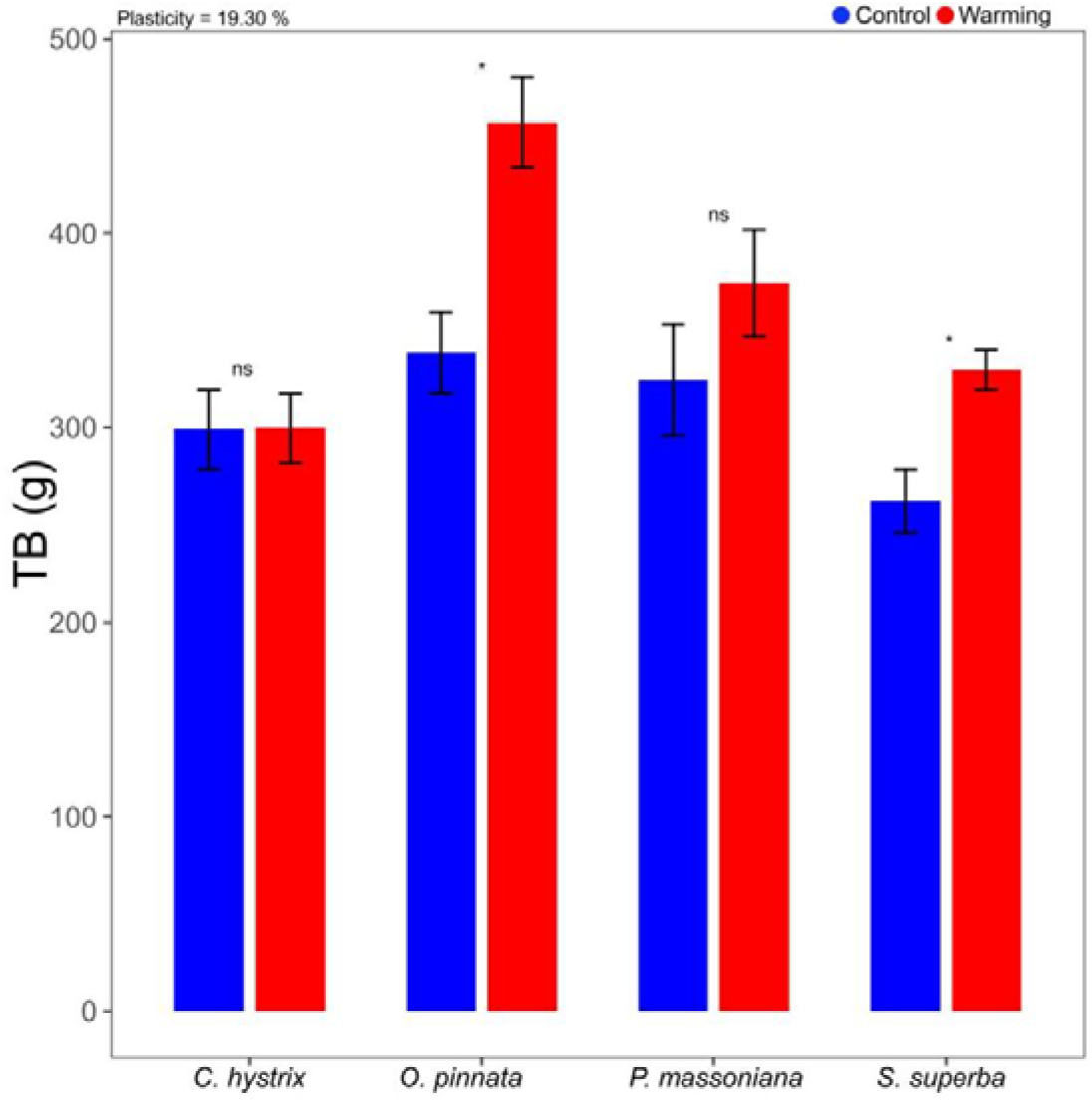
Total biomass of *Castanopsis hystrix*, *Ormosia Pinnata, Pinus massoniana and Schima superba* grown in the warming site and control site. Error bars are standard error (n= 5). Asterisks (*), (**) and (***) indicate that there are significant differences at *p*< 0.05, *p*< 0.01 and *p*< 0.001 between the warming and the control plots, respectively. Mean values of plasticity of the four species are reported for each trait.

**Fig. S2.**
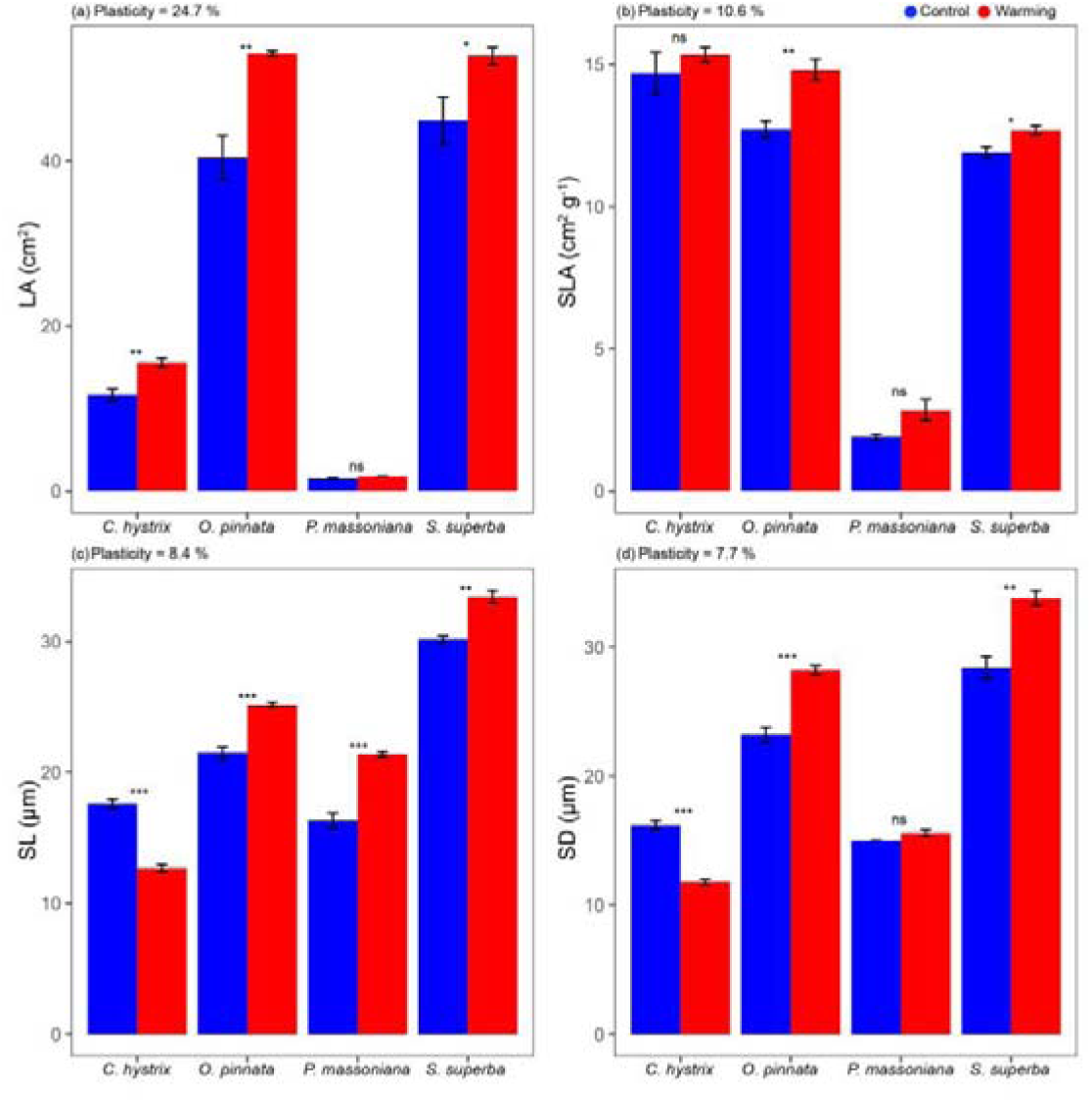
Leaf area (a), specific leaf area (b), stomatal length (c) and stomatal density (d) for *Castanopsis hystrix*, *Ormosia Pinnata, Pinus massoniana and Schima superba* grown in the warming site and control site. Error bars are standard error (n= 5). Asterisks (*), (**) and (***) indicate that there are significant differences at *p* < 0.05, *p* < 0.01 and *p* < 0.001 between the warming and the control plots, respectively. Mean values of plasticity of the four species are reported for each trait.

**Fig. S3.**
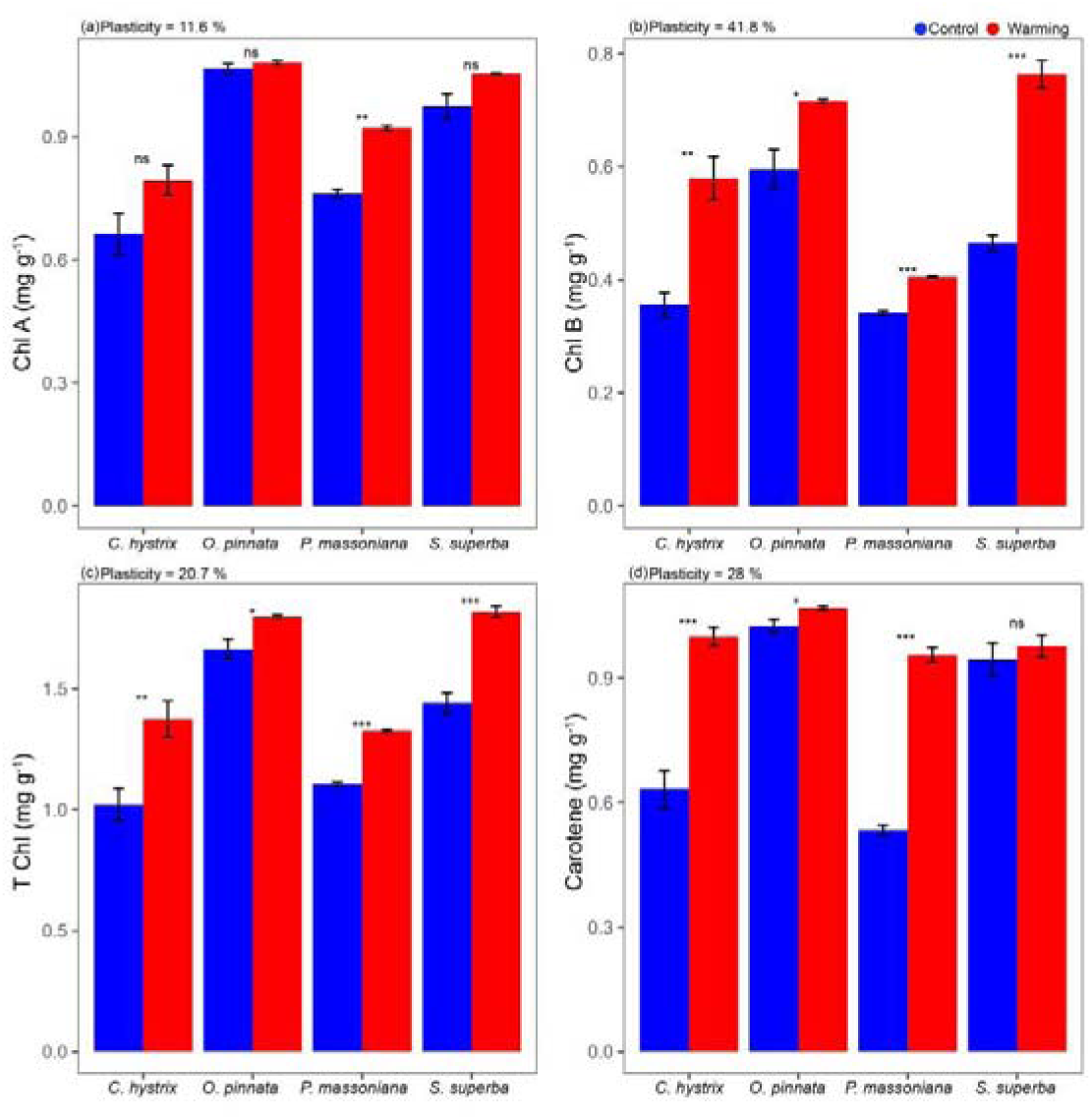
CHL A (a), CHL B (b), T Chl (c) and Carotenoids (d) for *Castanopsis hystrix*, *Ormosia Pinnata, Pinus massoniana and Schima superba* grown in the warming site and control site. Error bars are standard error (n= 5 trees). Asterisks (*), (**) and (***) indicate that there are significant differences at *p*< 0.05, *p*< 0.01 and *p*< 0.001 between the warming and the control plots, respectively. Mean values of plasticity of the four species are reported for each trait.

**Fig. S4.**
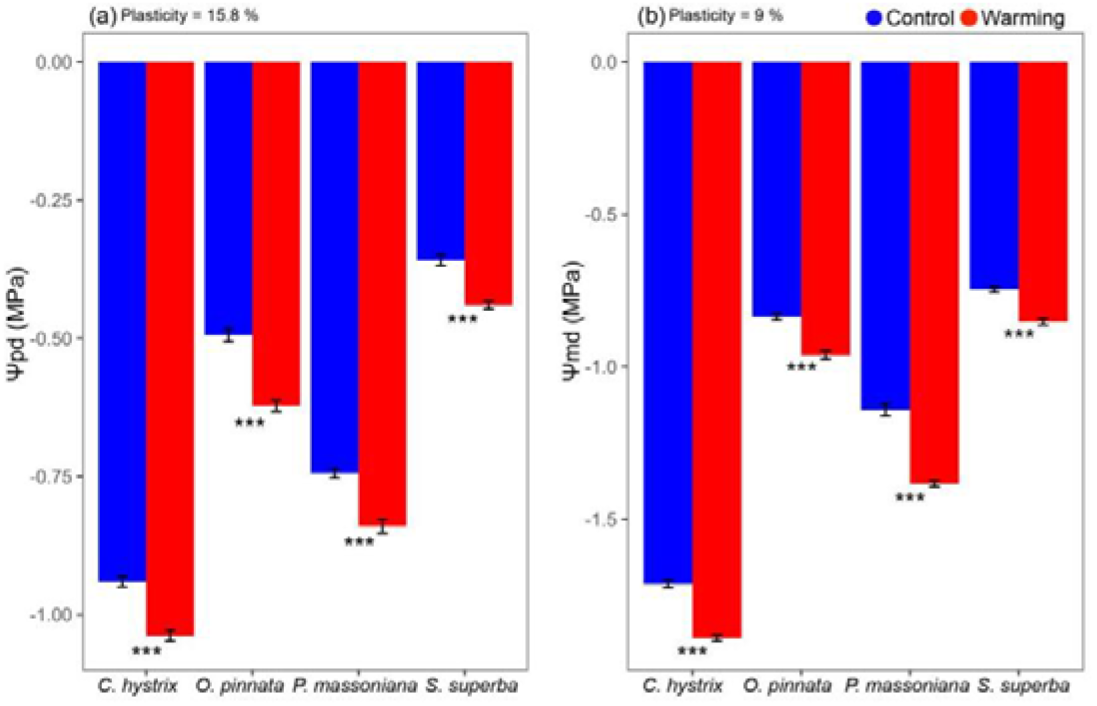
Ψpd (a) and Ψmd (b) for *Castanopsis hystrix*, *Ormosia Pinnata, Pinus massoniana and Schima superba* grown in the warming site and control site. Error bars are standard error (n= 5). Asterisks (*), (**) and (***) indicate that there are significant differences at *p*< 0.05, *p*< 0.01 and *p*< 0.001 between the warming and the control plots, respectively. Mean values of plasticity of the four species are reported for each trait.

**Fig. S5.**
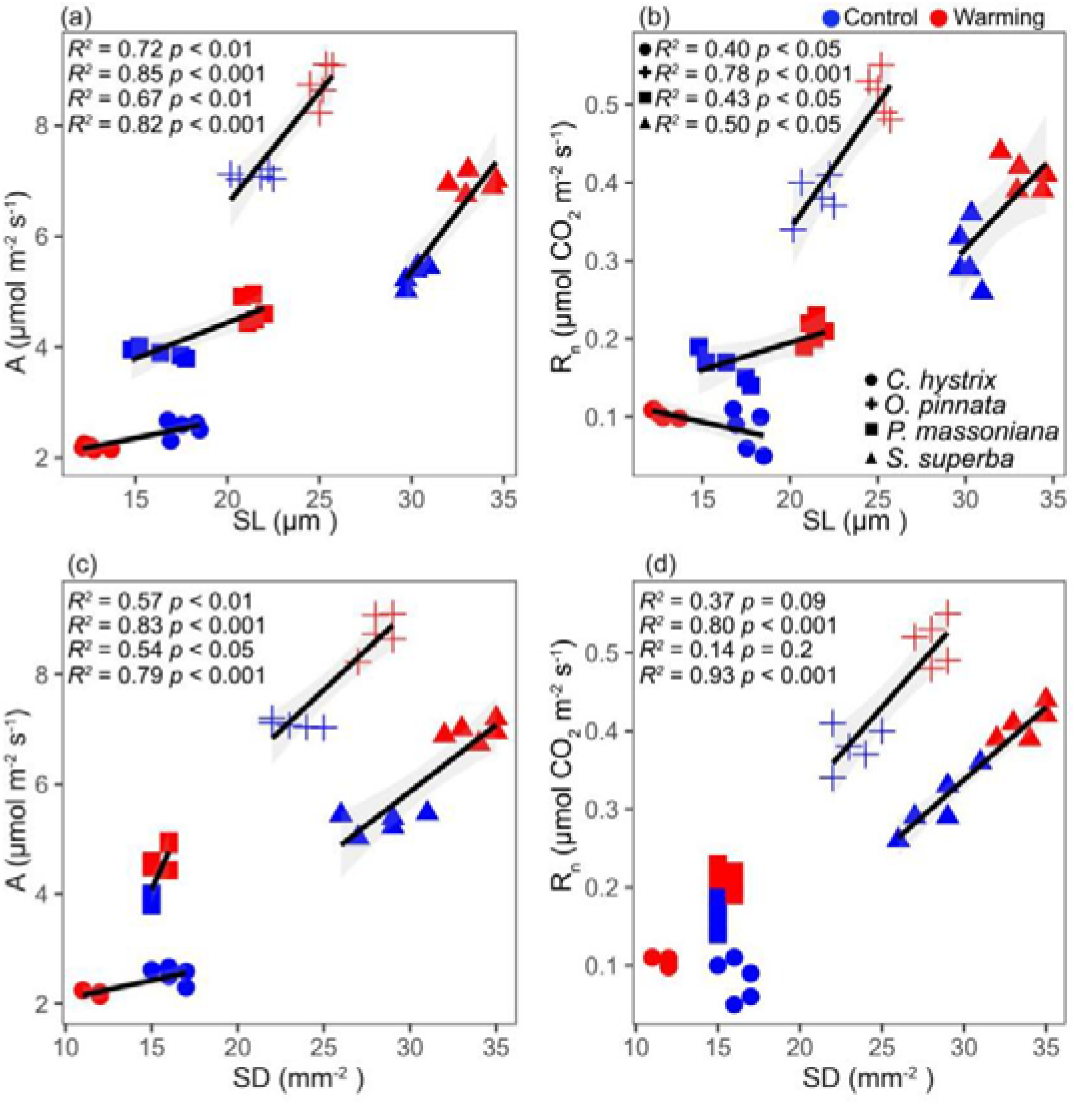
The relationships of photosynthetic rate A with SL (a) and SD (c) and night respiration (R_n_) with SL (b) and SD (d). Species under control are shown in blue; Warming condition species in red. Only statistically significant relationships (*p*<D0.05) are shown as solid lines with shaded areas representing the 95% confidence interval.

**Fig. S6.**
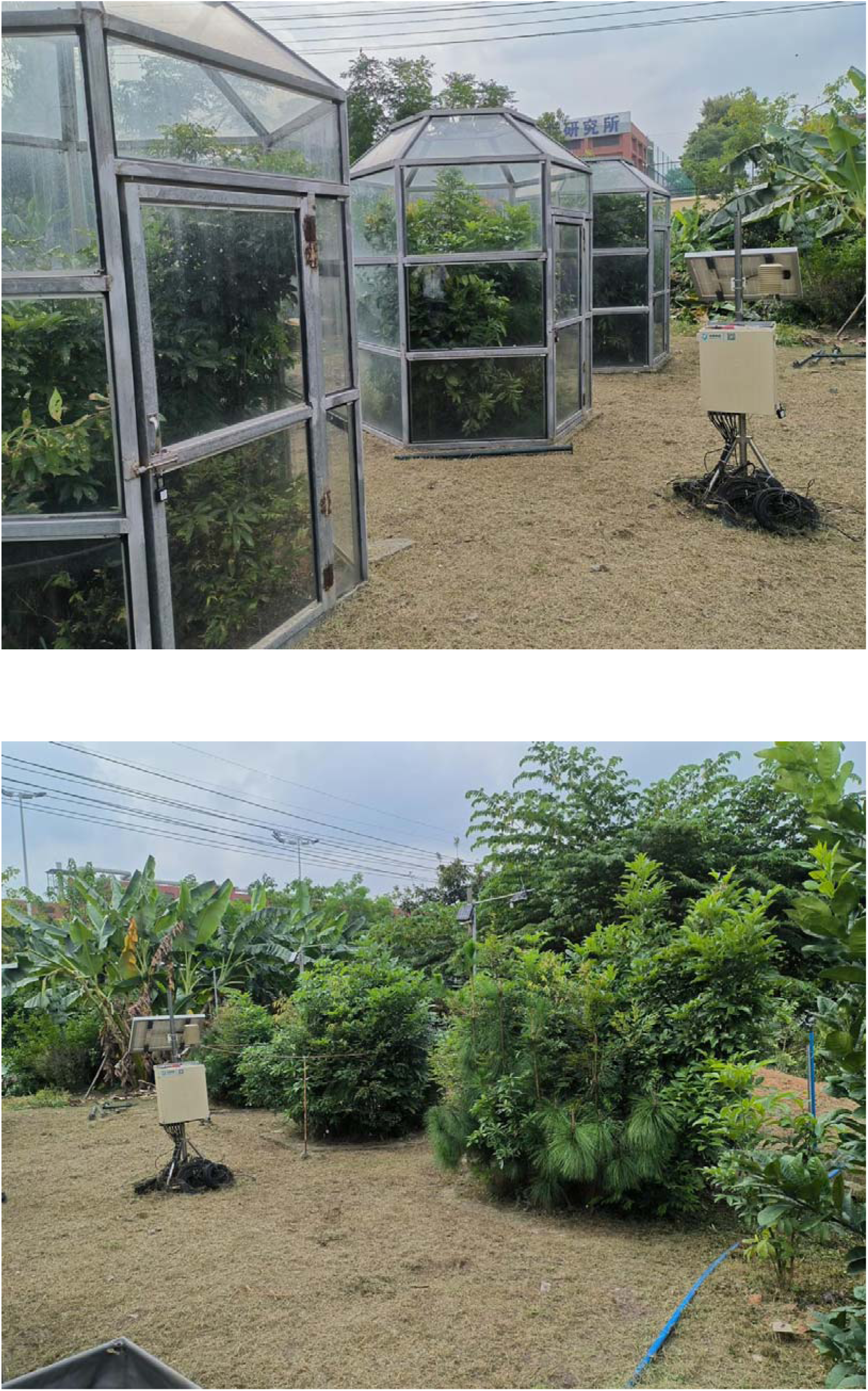
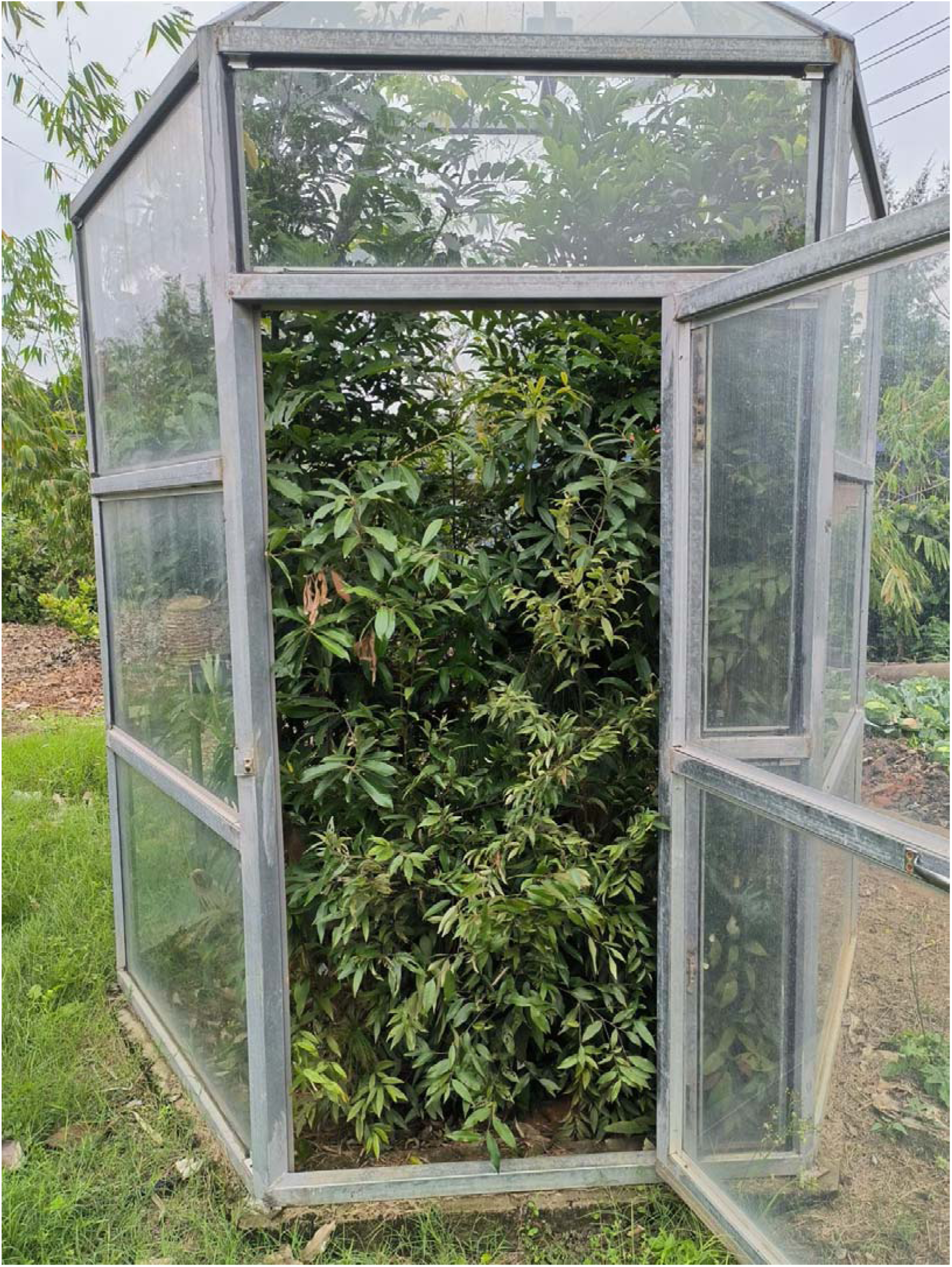
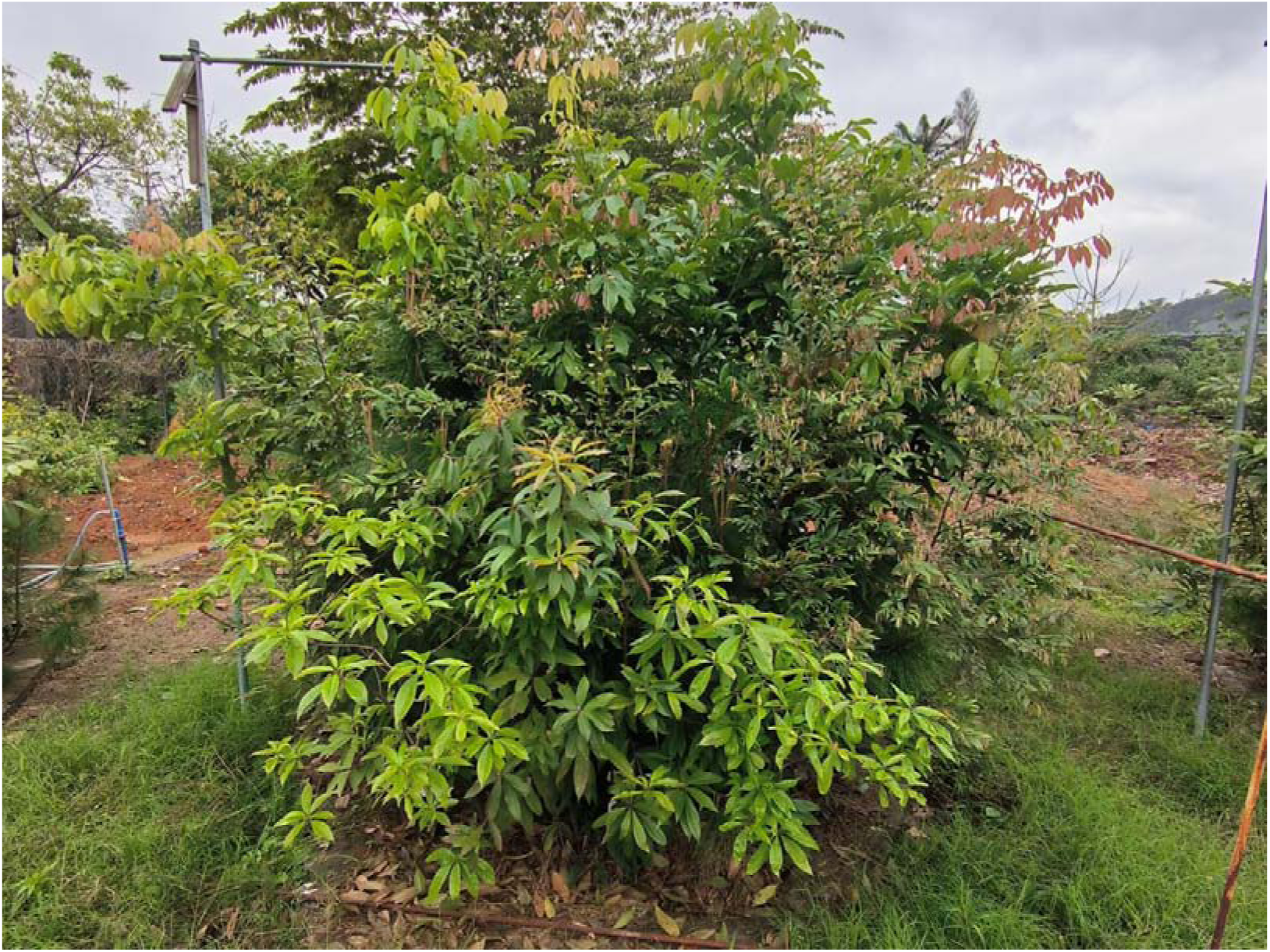
Field experimental design at South China Botanical Garden, China, showing (a) control plots under ambient conditions and (b) warming plots enclosed in open-top chambers (OTCs) to simulate increased temperature.

